# JuSpace: A tool for spatial correlation analyses of magnetic resonance imaging data with nuclear imaging derived neurotransmitter maps

**DOI:** 10.1101/2020.04.17.046300

**Authors:** Juergen Dukart, Stefan Holiga, Michael Rullmann, Rupert Lanzenberger, Peter C.T. Hawkins, Mitul A. Mehta, Swen Hesse, Henryk Barthel, Osama Sabri, Robert Jech, Simon B. Eickhoff

## Abstract

Recent studies have shown that drug-induced spatial alteration patterns in resting state functional activity as measured using magnetic resonance imaging (rsfMRI) are associated with the distribution of specific receptor systems targeted by respective compounds. Based on this approach, we introduce a toolbox (JuSpace) allowing for cross-modal correlation of MRI-based measures with nuclear imaging derived estimates covering various neurotransmitter systems including dopaminergic, serotonergic, noradrenergic, and GABAergic (gamma-aminobutric acid) neurotransmission. We apply JuSpace to two datasets covering Parkinson’s disease patients (PD) and risperidone-induced changes in rsfMRI and cerebral blood flow (CBF). Consistently with the predominant neurodegeneration of dopaminergic and serotonergic system in PD, we find significant spatial associations between rsfMRI activity alterations in PD and dopaminergic (D2) and serotonergic systems (5-HT1b). Risperidone induced CBF alterations were correlated with its main targets in serotonergic and dopaminergic systems. JuSpace provides a biologically meaningful framework for linking neuroimaging to underlying neurotransmitter information.

## Introduction

Magnetic resonance imaging (MRI)- derived measures are now commonly applied to study brain function and structure in health and disease (Good et al., 2001; Bohanna et al., 2008; Bloudek et al., 2011; Drysdale et al., 2017). Voxel- and region-wise analyses are commonly applied to study associations between task-based (tbfMRI) and resting state (rsfMRI) MRI measures and observed symptoms, behaviour or genetic information (Meyer-Lindenberg and Weinberger, 2006; Thompson et al., 2014; van Erp et al., 2015). RsfMRI measures provide replicable pathophysiological marker for various behaviours as well as different neurological and psychiatric conditions (Bernhardt et al., 2010; Telesford et al., 2010; Holiga et al., 2019). Despite this valuable information, biological and methodological limitations are imposed with respect to interpretation of the outcomes of voxel- and region-wise analyses.

From a methodological point of view, analyses of tbfMRI and rsfMRI are often limited by the rather low to moderate test-retest reliability of the commonly applied voxel- and atlas-based measures (Holiga et al., 2018). This low test-retest reliability and the resulting low signal-to-noise ratios impede constraints on the ability of fMRI to identify robust and replicable associations. Correspondingly, to date both tbfMRI and rsfMRI failed to achieve integration into routine clinical applications for the suggested indications (Lee et al., 2013; Leuthardt et al., 2018). Recent studies have shown that the overall spatial activity patterns (i.e. the relative within-subject activation of one region over another) of both tbfMRI and rsfMRI measures provide much more reliable marker as compared to standard voxel- and region-wise analyses (Dukart et al., 2018; Holiga et al., 2018). Making use of this higher reliability may therefore represent a viable way of improving the replicability of fMRI applications.

From biology point of view, standard analyses of fMRI data focus on identification of voxel- or region-wise signals associated with a specific condition. Whilst providing information about the spatial location of respective signals, such analyses do not allow drawing conclusions on potential neurophysiological mechanisms underlying the observed associations. To overcome this limitation, several recently published studies made use of spatial associations between underlying biology and observed imaging alterations by correlating *ex vivo* micro RNA spatial expression patterns with different imaging measures (Rizzo et al., 2016; Selvaggi et al., 2018; Liu et al., 2019). The major idea behind such analyses is that disease- or drug-induced changes in imaging measures occur in association with availability of a specific tissue property (i.e. expression of a specific receptor) that is affected by the respective condition. For example, in a disease that is primarily associated with loss of dopaminergic neurons, one would expect strongest imaging changes in regions, which contain many of such neurons in healthy individuals. Whilst promising, this approach also makes several assumptions that do not necessarily hold or are unknown for many of the underlying systems (Unterholzner et al., 2020). For example, correlations with mRNA expression imply that the respective genes are transcribed and lead to measurable changes of tissue structure or function. A viable way of making use of this concept whilst avoiding these assumptions is by integration of positron emission tomography (PET) or single photon computed emission tomography (SPECT) derived tissue property maps. Recent advancements in PET and SPECT tracer development resulted in a variety of novel tracers that can reliably measure the availability of specific receptors but also functional aspects such as synthesis capacity across a variety of neurotransmitters (Smith et al., 1998; Mawlawi et al., 2001; McCann et al., 2005; Lehto et al., 2015; Beliveau et al., 2016). Such PET- and SPECT- derived maps provide a more direct measurement of specific tissue properties as compared to mRNA expression. In line with that, we have shown that MRI-derived spatial activity patterns induced by different drugs correlate with PET- and SPECT- derived information that are associated with the mechanism of action of respective compounds (Dukart et al., 2018).

Here we introduce the JuSpace toolbox allowing for spatial correlation of MRI-based or other imaging modalities with PET- and SPECT- derived maps covering a variety of neurotransmitter systems. To demonstrate its utility, we deploy the toolbox to rsfMRI data of Parkinson’s disease (PD) patients on and off levodopa – a disease with devastating effects on multiple neurotransmitter systems, including major contributions from dopamine and serotonin (Booij et al., 1997; Pagano et al., 2017) – as well as to cerebral blood flow data of healthy volunteers scanned on and off risperidone – an antipsychotic with a serotonergic and dopaminergic mechanism of action.

## Methods

### Software description

The main idea of JuSpace is to test if MRI-derived information is spatially structured in a way that reflects the distribution of specific biologically interpretable tissue properties as derived from PET and SPECT modalities. JuSpace is a comprehensive license-free toolbox (only for non-commercial use) for the integration of PET- and SPECT- derived modalities with other brain imaging data. However, we do ask to cite the specific references for the PET and SPECT maps, which are used for the respective analyses. The references are provided in Table 1. The currently released version is available at https://github.com/juryxy/JuSpace). JuSpace has been developed in the Matlab environment (Matlab 2017a or higher) and requires Statistical Parametric Mapping Software (SPM12, https://www.fil.ion.ucl.ac.uk/spm/software/spm12/) as well as the Matlab Statistics toolbox to be installed. It consists of a group of Matlab functions together with PET receptor maps covering various receptor systems. JuSpace provides a graphical user interface (Figure 1a) as well a direct call to the respective functions. There are no specific system requirements.

**Table 1.**
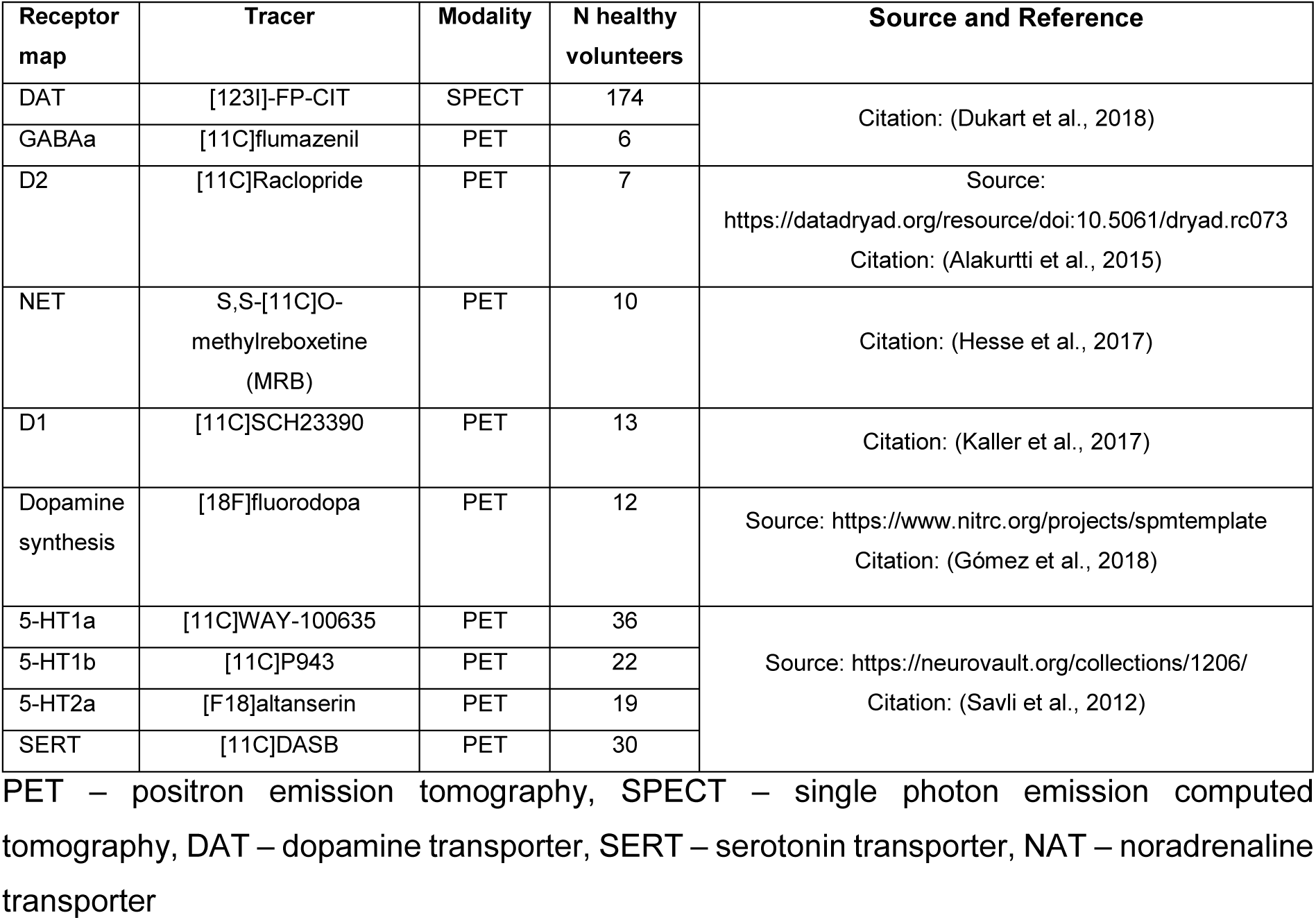
Receptor maps included in the JuSpace toolbox

| Receptor map | Tracer | Modality | N healthy volunteers | Source and Reference |
| --- | --- | --- | --- | --- |
| DAT | [123I]-FP-CIT | SPECT | 174 | Citation: (Dukart et al., 2018) |
| GABAA | [11C]flumazenil | PET | 6 |  |
| D2 | [11C]Raclopride | PET | 7 | Source:<br>https://datadryad.org/resource/doi:10.5061/dryad.rc073<br>Citation: (Alakurtti et al., 2015) |
| NET | S,S-[11C]O-methylreboxetine (MRB) | PET | 10 | Citation: (Hesse et al., 2017) |
| D1 | [11C]SCH23390 | PET | 13 | Citation: (Kaller et al., 2017) |
| Dopamine synthesis | [18F]fluorodopa | PET | 12 | Source: https://www.nitrc.org/projects/spmtemplate<br>Citation: (Gómez et al., 2018) |
| 5-HT1a | [11C]WAY-100635 | PET | 36 | Source: https://neurovault.org/collections/1206/<br>Citation: (Savli et al., 2012) |
| 5-HT1b | [11C]P943 | PET | 22 |  |
| 5-HT2a | [F18]altanserin | PET | 19 |  |
| SERT | [11C]DASB | PET | 30 |  |

PET – positron emission tomography, SPECT – single photon emission computed tomography, DAT – dopamine transporter, SERT – serotonin transporter, NAT – noradrenaline transporter

**Figure 1.**
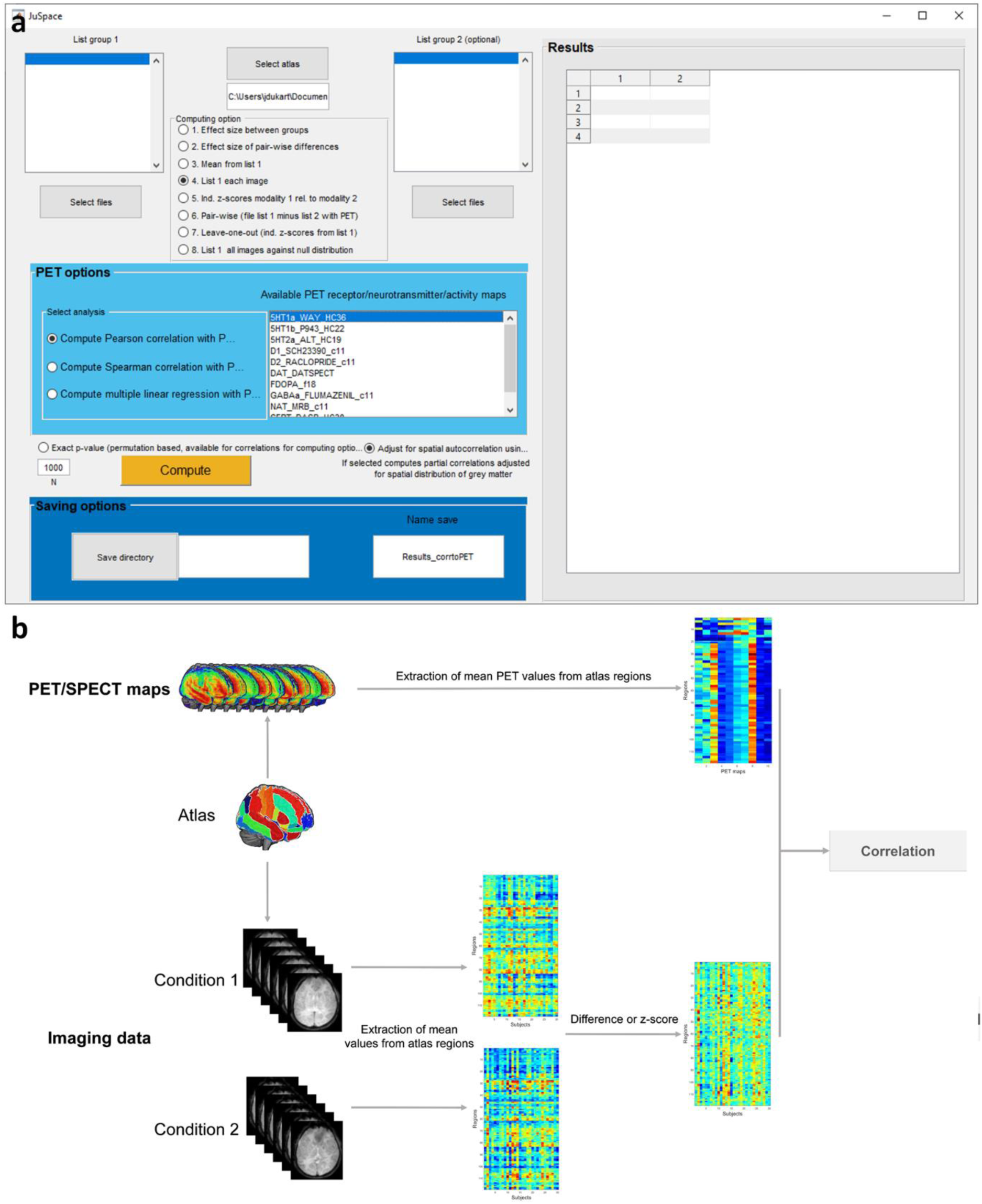
JuSpace toolbox, a) User interface of the JuSpace toolbox, b) Schematic work flow of the JuSpace toolbox

### Included PET maps

All PET and SPECT maps are free for non-commercial distribution and were previously published as described in Table 1 and in the release notes provided with the toolbox. All PET maps were derived from average group maps of different healthy volunteers and linearly rescaled to a minimum of 0 and a maximum of 100:

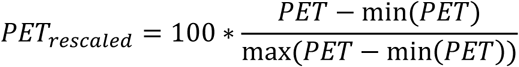

PET maps covering the following receptor types are included in the first release: 5-HT1a (serotonin 5-hydroxytryptamine receptor subtype 1a), 5-HT1b (5-HT subtype 1b), 5-HT2a (5- HT subtype 2a), D1 (dopamine D1), D2 (dopamine D2), DAT (dopamine transporter), F-DOPA (dopamine synthesis capacity), GABAa (gamma-aminobutric acid), NAT (noradrenaline transporter) and SERT (serotonin transporter) (for references see Table 1).

### Workflow

The analysis workflow starts with the user selecting the imaging (i.e. MRI) data to correlate with provided PET and SPECT maps. Either data for a single modality (“files 1” only) or data to generate a contrast between conditions (“files 1” and “files 2”, i.e. patients vs. healthy controls or pre- vs. post-treatment data) are entered as input. The default atlas is the neuromorphometrics atlas from SPM12 (Friston et al., 1994) excluding all white matter and cerebrospinal fluid regions. A symmetric version of the atlas with bilateral regions of interest (left side flipped) is also included in the release. Neuromorphometrics atlas probability tissue labels were derived from the “MICCAI 2012 Grand Challenge and Workshop on Multi-Atlas Labeling” (https://masi.vuse.vanderbilt.edu/workshop2012/index.php/Challenge_Details). The atlas can be changed to any custom atlas using the “Select atlas” button. The atlas is used to extract mean regional values from the entered MRI modalities to be correlated with respective values from selected PET and SPECT maps. An atlas is needed as correlation of voxel-wise maps would result in highly inflated degrees of freedom. The number of distinct spatial features strongly depends on data smoothness but is typically in the range of several hundred or more distinct resolution elements (Mikl et al., 2008). In that sense, the default atlas with 119 regions provides a conservative estimate for the effective degrees of freedom. Next, the computing option is selected. Currently available options are:

1. Effect size between groups (computes Cohen’s d for each atlas region between files selected in files 1 and files 2)
2. Effect size of pair-wise differences (computes Cohen’s d for pair-wise differences between files in files 1 relative to files 2)
3. Mean from files 1 (computes mean value per atlas region for files 1)
4. Files 1 each image (extracts mean value per atlas region for each file from files 1)
5. Compute individual z-score maps for each file in files 1 relative to files 2
6. Computes pair-wise differences between files 1 and files 2
7. Computes leave-one-out z-scores maps for each file in files 1 relative to other files in files 1
8. Extracts mean value per atlas region for each file from files 1 and compares correlation coefficients for all images against null distribution

Further, the analysis type is selected (Pearson correlation, Spearman correlation or multiple linear regression). PET and SPECT maps can be selected by clicking on the respective name (multiple selection is supported, i.e. hold control button during selection on a Windows machine). The currently available PET and SPECT maps are listed in Table 1.

Further, the option for exact permutation based p-value (only for computing options (1),(2),(5) and (6)) can be selected (as described below). Additionally, the option is provided to adjust for spatial autocorrelation using the grey matter probability map TPM.nii from SPM12. The saving directory has to be specified using the “Save directory” button. The “Compute” button initiates the computation by calling the function “compute_DomainGauges” with the chosen computational parameters.

### Computational workflow

All provided files as well as the selected PET maps are loaded into the atlas space as mean value per file and region (Figure 1b). Depending on the choice of the computing option, a spatial correlation or multiple linear regression is then computed between the selected PET maps and the extracted values as per selected computing option. In case adjustment the optional adjustment for spatial autocorrelation was selected, a partial spatial correlation is computed between both adjusting for local grey matter probabilities as estimated from TPM.nii provided with SPM12. In case of multiple linear regression, local grey matter probabilities are added into the model. For correlation analyses, Fisher’s z-transformed coefficients are provided as well as the original correlation coefficients. The distribution of Fisher’s z transformed correlation coefficients or regression coefficients for computing options (5)-(8) is compared against null distribution using one-sample t-tests. For group-level computing options (1)-(4), the p-value is provided directly for the specific correlation /multiple linear regression analysis.

### Permutation statistics

An optional exact orthogonal permutation based p-value can be computed for correlational analyses (Pearson and Spearman) for computing options (1),(2),(5) and (6). The orthogonal permutation approach ensures that the shuffled labels vector is uncorrelated with the initial label vector providing a more valid null distribution (Aickin, 2010). The exact p-value is then computed using add-one discounting. For the within-subject designs (computing option (2) and (6)), the permutations are performed by random switching of 50% of the data between files 1 and files 2 whilst maintaining pairwise associations. For the between-subject designs (computing options (1) and (5)), the files from both groups are randomly permuted across files 1 and files 2 whilst maintaining the initial relative ratios of both groups in each permutation. For options (5) and (6), the exact p-value is computed as the number of mean absolute correlation coefficients across permutations exceeding the observed mean absolute correlation coefficient.

### Correlation between PET and SPECT maps

We computed Spearman correlations using computing option (4) with and without adjustment for auto-correlation to understand the interdependencies between all included PET and SPECT maps. Significant positive correlations were observed between most PET maps with strongest correlations of up to rho=.89 for GABAa and 5-HT2a and rho=.88 for DAT and SERT (all p<.001) (Figure 2). The only significant but weak negative correlation was observed between 5-HT1a and D2 (rho=-0.19, p=.037). To illustrate the utility of JuSpace we applied it to two datasets capturing disease- and drug-induced activity alterations as measured using rsfMRI.

**Figure 2.**
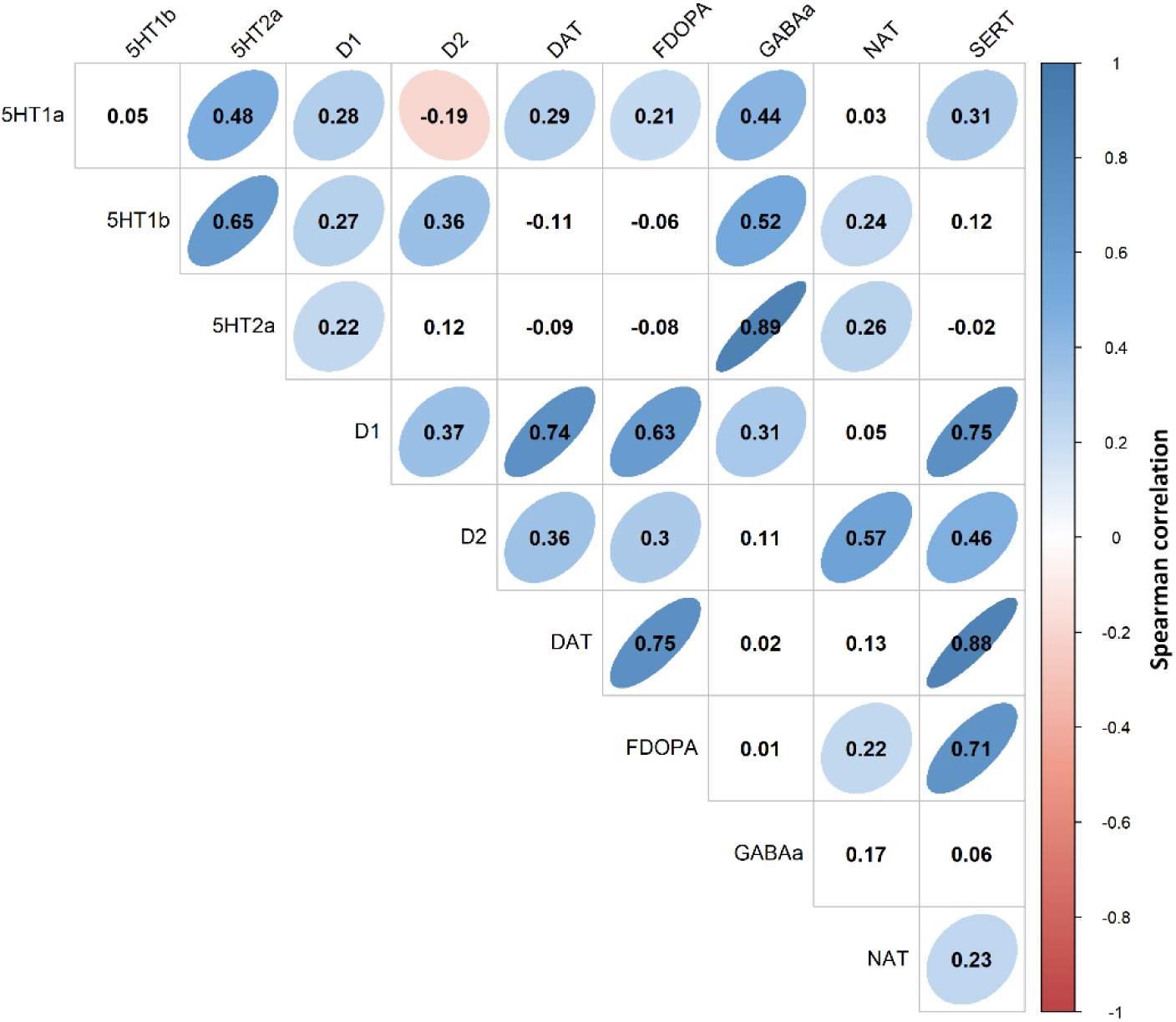
Results of spatial correlation analyses between PET and SPECT derived neurotransmitter maps. The displayed numbers are the observed Spearman correlation coefficients. Significant correlations are highlighted by underlying ellipses. DAT – dopamine transporter, SERT – serotonin transporter, NAT – noradrenaline transporter, FDOPA – Fluorodopa

### Application example 1

#### Dataset

To demonstrate the functionality of the JuSpace we applied it to an rsfMRI dataset of 30 PD patients scanned on and off levodopa as compared to 30 age and sex matched healthy controls (HC). A detailed description of the dataset as well of image acquisition and preprocessing is provided in Supplement 1. Fractional Amplitude of Low Frequency Fluctuations (fALFF) was computed as a measurement of local activity using the REST toolbox with default parameters (linear detrending and 0.01-0.08 Hz band-pass filtering).

#### Spatial correlation analyses

We aimed to evaluate if fALFF alterations in PD patients on and off levodopa (individual z-scores) as compared to HC are correlated with specific neurotransmitter systems. For this, we used the JuSpace toolbox to compute Spearman correlation coefficients between respective measures (computing option (5)) and the above PET maps included in the toolbox. We further aimed to test for the effects of levodopa on the fALFF maps. For this, we correlated the single subject pairwise differences between the fALFF maps with and without levodopa with the above PET maps (computing option (6)). Exact permutation-based p-values (with 10000 permutations) were computed for all analyses to test if the mean correlation coefficients observed across subjects are significantly different from the null distribution. All analyses were false discovery rate (FDR) corrected for the number of tests for each group comparison.

#### Voxel-wise analyses

To compare the sensitivity and the information provided by the spatial correlation approach we additionally performed standard voxel-wise analyses in SPM12 comparing either PD patients on and off levodopa to HC (two-sample t-tests, including age and sex as covariates), or PD patients on levodopa to their off levodopa state (paired t-tests). The focus of the above spatial correlation analyses was on evaluating similarity of PD-related and drug-induced spatial patterns with specific PET maps. To visualize all regions showing strongest respective changes we applied a liberal voxel-wise threshold of p<.05 combined with a cluster threshold of 100 voxels. Additionally, we report all contrasts which survive classical voxel-wise whole-brain family-wise error correction (p<.05 FWE corrected) for multiple comparisons.

### Application example 2

#### Dataset and image processing

In a second application example, we applied JuSpace to a CBF dataset of healthy volunteers (N=21) scanned on placebo and on a low and high dose of the dopamine antagonist risperidone (0.5 and 2 mg) – a serotonin and dopamine antagonist. The risperidone cohort is described in detail in Supplement 1 and in a previous publication (Hawkins et al., 2017). Pre- processing of the CBF risperidone data is described elsewhere (Dukart et al., 2018). In brief, it comprised computation of quantitative CBF maps, normalization into MNI space using structural information and masking of non-grey matter voxels. Additionally, smoothing with Gaussian kernel of 8 mm FWHM was applied prior to voxel-wise analyses.

#### Spatial correlation analyses

For the risperidone dataset, we tested for the effects of high and low dose of risperidone as compared to placebo and to each other by computing Spearman correlation coefficients between respective within-subject pairwise differences (computing options (6)). Exact permutation-based p-values (10000 permutations) were computed for all analyses. All analyses were FDR corrected according to the number of tests for each group comparison.

#### Voxel-wise analyses

We computed a within subject ANOVA to compare risperidone high dose, risperidone low dose and placebo conditions using pair-wise t-contrasts. Same voxel- and cluster-wise thresholds as for the PD dataset were applied.

## Results

### Application example 1

#### Results of spatial correlation analyses

Individual fALFF alterations in PD patients off levodopa as compared to HC were significantly associated with spatial distribution of D2 (p<.001) and 5-HT1b (p=.003) receptors as derived from healthy subjects (Figure 3a). Similarly, fALFF alterations in PD patients on levodopa were significantly associated with availability of D2 (p=.002) and 5-HT1b receptors (both p=.008) (Figure 3b). There was no significant difference between levodopa on and off conditions (all p>.29) (Figure 3c).

**Figure 3.**
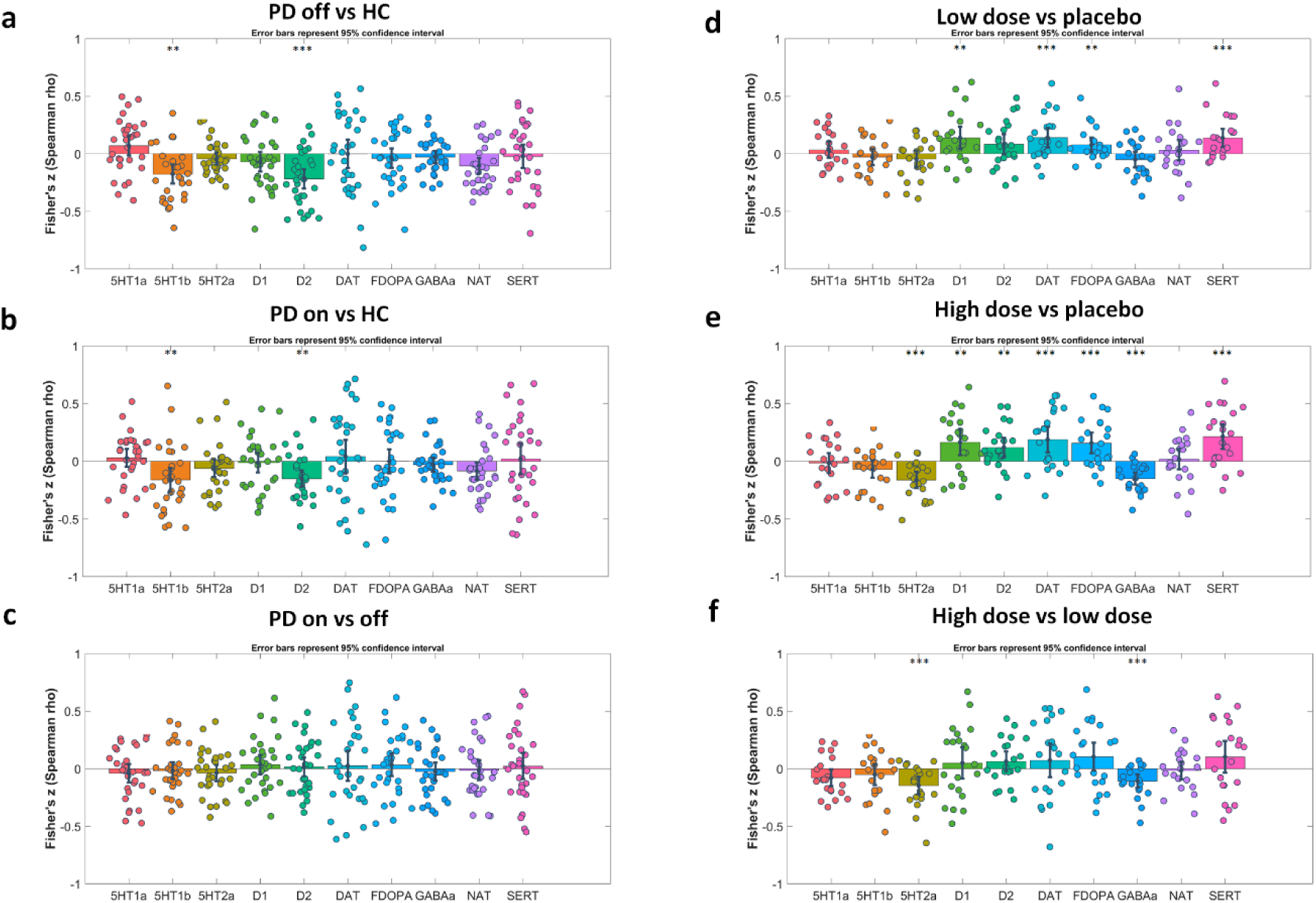
Results of spatial correlation analyses for PD and risperidone datasets. PD – Parkinson’s disease, HC – healthy controls, DAT – dopamine transporter, SERT – serotonin transporter, NAT – noradrenaline transporter, FDOPA – Fluorodopa

#### Results of voxel-wise analyses

In voxel-wise comparisons to HC, decreased fALFF was observed in PD on and off levodopa conditions in an extensive network covering predominantly prefrontal, parietal, cerebellar, basal ganglia, supplementary and primary motor regions (Figure 4a, b). Increased fALFF was observed primarily in temporal and orbitofrontal cortices. Decreased fALFF was observed in PD on levodopa as compared off levodopa in prefrontal, left temporal and right parietal cortices (Figure 4c). None of the effects survived whole-brain voxel-wise correction for multiple comparisons

**Figure 4.**
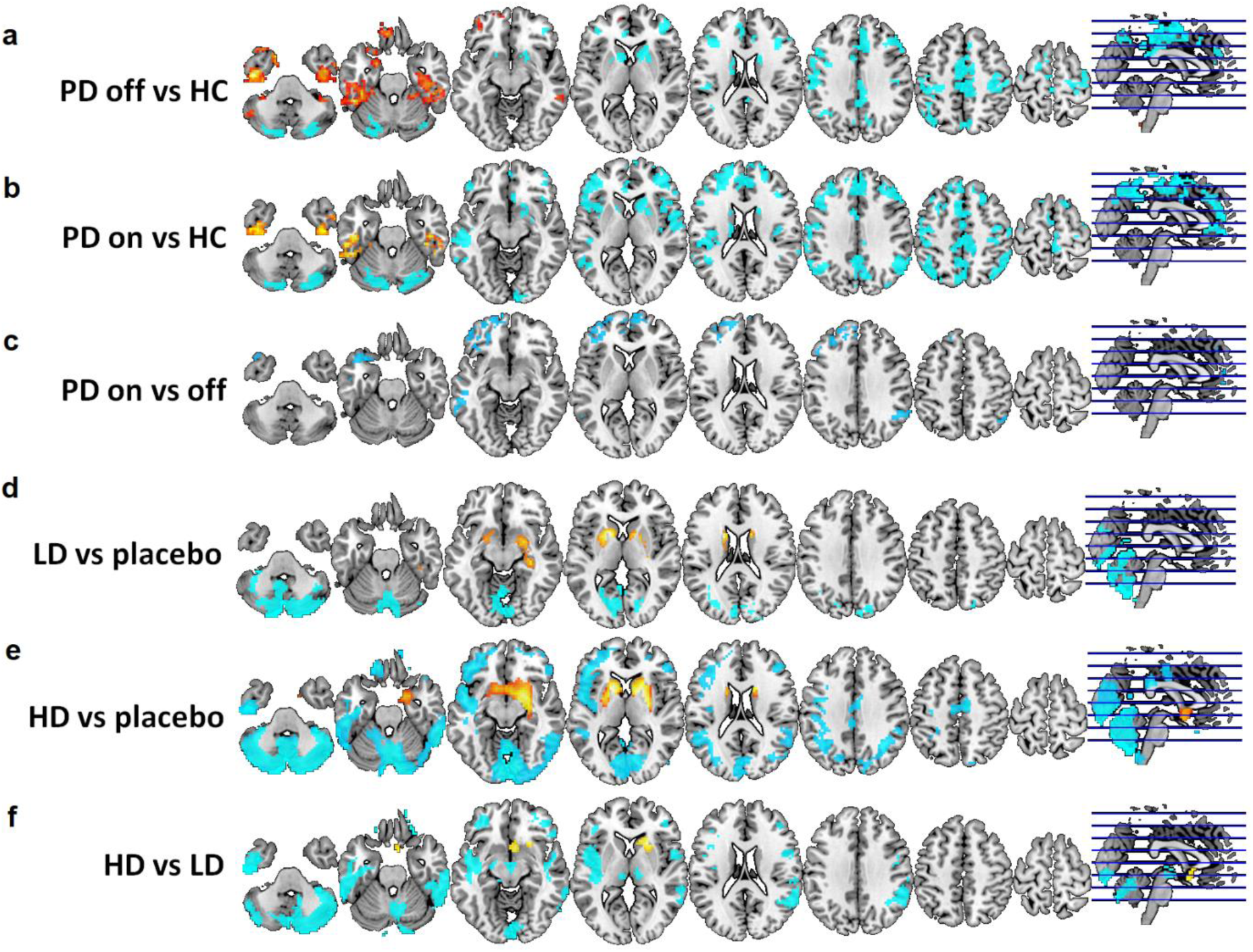
Results of voxel-wise analyses for PD and risperidone datasets. Orange and red colors indicate increased fALFF (for PD) or CBF (for risperidone) in the first mentioned condition/group. Cyan colors indicate decreased fALFF (for PD) or CBF (for risperidone) in the first mentioned condition/group. PD – Parkinson’s disease, HC – healthy controls, HD – high dose of risperidone, LD – low dose of risperidone,

### Application example 2

#### Results of spatial correlation analyses

CBF alterations induced by the low dose of risperidone as compared to placebo were significantly associated with D1 (p=.002), DAT (p<.001), F-Dopa (p=.006) and SERT (p<.001) maps (Figure 3d). CBF changes induced by the high dose of risperidone were significantly correlated with 5-HT2a (p<.001), D1 (p=.003), D2 (p=.003), DAT (p<.001), GABAa (p<.001) and SERT (p<.001) maps (Figure 3e). CBF changes with the high dose as compared to low dose of risperidone were significantly associated with 5-HT2a and GABAa receptor availability (both p<.001) (Figure 3f).

#### Results of voxel-wise analyses

Increased CBF was observed in basal ganglia comparing high and low dose of risperidone to placebo (Figure 3d,e). Reduced CBF in low dose as compared to placebo was predominantly restricted to occipital and cerebellar cortices. High dose associated changes were more extensive covering prefrontal, posterior cingulate, cerebellar temporal, parietal and occipital regions. High dose as compared to low dose of risperidone showed an increase in CBF in the right corpus striatum (Figure 3f). Reduced CBF was observed with high dose in prefrontal, temporal, occipital, parietal and cerebellar regions. Only the reduced CBF in cerebellar and occipital regions in high dose as compared to placebo survived whole-brain voxel-wise correction for multiple comparisons.

## Discussion

Here we introduce the JuSpace Toolbox, an integrated system for the comparison of PET and SPECT derived neurotransmitter maps with other imaging modalities such as rsfMRI data. The software tests for associations between the imaging data of interest and a list of included PET and SPECT maps by computing correlations or multiple linear regressions.

JuSpace allows for an easy integration of neuroimaging data with PET-derived receptor maps. JuSpace is a user-friendly tool allowing user interface-based applications by researchers with limited programming experience as well as direct function calls. The choice of settings for the analyses is kept to a minimum. The toolbox further supports an easy integration of other atlases and other PET-derived information, provided they have the same resolution, by simply adding the respective maps in MNI space into the PET atlas directory. JuSpace is designed to test the hypotheses that the spatial structure of imaging alterations induced i.e. by disease or drug is associated with the availability of a specific receptor across the brain. Besides direct correlation between the imaging data of interest and the available PET maps, the software supports simple between- and within-subject designs by computing effect sizes, z-scores and pair-wise differences. For both, between- and within-subject designs the toolbox provides more rigorous permutation based statistics. Usage of these exact statistics for both options is strongly recommended to avoid erroneous assumptions on data distribution or actual spatial degrees of freedom.

As compared to available imaging-genomic toolboxes correlating imaging information with the spatial information derived from few donors from the Allen Brain Atlas of post-mortem mRNA expression JuSpace carries the advantage of making less assumptions about underlying biology (i.e. unknown transcription of respective mRNA into specific tissue properties) (Rizzo et al., 2016). It also allows for evaluation of spatial associations with PET-derived transmitter synthesis information that are only available from in vivo studies, i.e. F-Dopa PET-derived dopamine synthesis capacity.

### Conclusions from the application examples

Here we applied the JuSpace toolbox to two datasets covering disease related rsfMRI alterations in PD as well drug-induced CBF alterations in healthy controls. We show that rsfMRI activity alterations in PD on and off levodopa are closely associated with availability of D2 and 5-HT1b receptors. These results are closely in line with the well-established affectedness of the dopaminergic and serotonergic systems in PD (Booij et al., 1997; Pagano et al., 2017). Furthermore, the consistency of the effects obtained with both scans in PD illustrates the robustness of findings obtained using spatial correlation analyses. We do not find significant associations between PET maps and fALFF differences between PD on and off levodopa. This is potentially explained by the effects being either subtle or not following the distribution of the specific receptor maps currently included in JuSpace.

Risperidone-induced brain activity alterations are associated with a variety of neurotransmitter systems including dopaminergic and serotonergic effects. In contrast to the highest affinity of risperidone to 5-HT2a followed by D2 the significant associations with the corresponding PET receptor maps only appeared with the high dose (Schotte et al., 1996). The strongest effects observed with low dose rather associate with DAT, SERT and D1 receptor maps. In our previous study (Selvaggi et al., 2018) we observed correlations with D2 receptors at both doses for the group averaged data. This study did not examine effects using other targets and did not evaluate individual differences. Importantly, the spatial correlation analysis relies on a direct translation of the effect observed on the specific receptor into the respective activity measurement, which may vary by drug and across subjects. The discrepancies observed suggest that the effects on different receptors may have different transfer function onto the observed CBF changes that is not directly associated with respective receptor affinities. This observation is in line with our previous work evaluating correlations with post-mortem receptor expression (Dukart et al., 2018). Overall, whilst the observed correlations with serotonergic and dopaminergic system are in line with the known mechanism of action of risperidone the correlation with GABAa receptor appearing with the high dose may appear unexpected as there is no reported affinity of risperidone to the respective receptor. There are two potential explanations for this effect. We observe a very strong positive correlation between GABAa and 5-HT2a receptor availability. In that sense, the observed correlation with both receptors is likely due to the collinearity of both receptor systems making it difficult to dissociate the specific effects on one of the two systems. Another possible explanation for the observed correlation with GABAa may be in the reported interdependence of both systems with 5-HT2a activation resulting in inhibition of GABAa currents (Feng et al. 2001). Similar to that, the strong correlation of PET maps may also explain the observed correlation of the low dose changes with DAT, SERT and D1 as all of the PET maps are strongly correlated. These strong cross-correlations between different PET maps indicate that caution is required with respect to interpretation of observed associations as being specific to a particular receptor. Further analyses adjusting for cross-correlation of the PET receptors such as the multiple linear regression analyses option also provided with the toolbox may facilitate the interpretation in such cases.

Whole-brain corrected analyses of both datasets only revealed differences between high dose of risperidone and placebo. The results applying a more liberal threshold were rather diffuse covering a wide-range of regions making it difficult to interpret the findings with respect to any specific anatomical circuitries. In contrast, spatial correlation analyses reveal a higher sensitivity to both PD- and risperidone-induced changes and additionally provide a biologically meaningful interpretation of the observed effects. Our results therefore suggest the higher sensitivity of the spatial correlation approach to detect disease- and drug-related activity alterations as compared to standard voxel-wise analyses. The likely reasons for the higher sensitivity of the spatial correlation as compared to voxel-wise analysis is the substantially higher reliability of spatial activity profiles as compared to classical voxel- or region-wise fALFF and CBF measures (Holiga et al., 2018).

Overall, the JuSpace toolbox allows for cross-modal evaluation of neuroimaging data alongside molecular imaging atlases. The inclusion of PET and SPECT atlases for different neurotransmitter systems allows for biologically meaningful evaluation and interpretation of the spatial patterns. This is a flexible platform enabling inclusion of user-defined atlases and other imaging modalities. As such, it has a great potential to improve and simplify multi-modal brain imaging research.

## Author’s contribution

JD wrote the manuscript. JD designed the overall study, wrote the toolbox performed all analyses and wrote the manuscript. MAM, PCTH and RJ contributed to the application examples. SH, MR, RL, SH, HB and OS contributed to creation of PET and SPECT maps. SBE contributed to the overall design of the study as well as to toolbox conceptualization. All authors reviewed and commented on the manuscript.

## Acknowledgments

The PD study was supported by the Czech Ministry of Health, grant AZV NV19-04-00233. This open source software code was developed in part or in whole in the Human Brain Project, funded from the European Union’s Horizon 2020 Framework Programme for Research and Innovation under the Specific Grant Agreement No. 785907 (Human Brain Project SGA2). The funders had no role in study design, data collection and analysis, decision to publish, or preparation of the manuscript.

## Software availability

The JuSpace toolbox is available at https://github.com/juryxy/JuSpace

