## Supplement 1 for "JuSpace: A tool for spatial correlation analyses of magnetic resonance imaging data with nuclear imaging derived neurotransmitter maps"

### Supplement 1, Dukart et al.

#### PD cohort

Thirty patients diagnosed with sporadic PD (PD duration 11.2±3.6 years) and 30 matched healthy controls were included in the study (Table S1). PD was diagnosed by neurologists specialized in movement disorders according to UK brain bank criteria [Hughes et al., 1992]. Other inclusion criteria were: PD onset after 45 years of age and Hoehn and Yahr stage from I to III. In addition, stable medication at least 4 weeks before enrollment, consisting of levodopa in monotherapy or in combination with dopamine agonists (pramipexole or ropinirole). Exclusion criteria were: actual or past psychotic symptoms, antipsychotic treatment, cognitive impairment (Montreal Cognitive Assessment –MoCA – below −1.5 SD compared to Czech norms), deep brain stimulation or jejunal levodopa infusion, any illness potentially affecting motor or cognitive state and any contraindication for MRI. Beside detailed clinical assessment for PD all subjects underwent an MRI assessment including acquisition of rsMRI and structural MRI data. Thereby, all HC underwent one MRI session and all PD patients two MRI sessions 14,7±11,1 days apart in OFF (UPDRS III: 30,8±9,8) and ON antiparkisonian medication (UPDRS III: 15,1±7,6) condition in a balanced cross-over design. Four days before OFF medication session, dopamine agonists were substituted by equivalent doses of levodopa in each patient (Tomlinson et al., 2010). Other anti-PD medications (selegiline, amantadine, anticholinergics) were suspended. While in the OFF state, the MRI was obtained after an overnight withdrawal of levodopa (at least 12 h), in the ON state, the MRI was obtained with optimal antiparkinsonian medication. All subjects gave their informed consent to participate in the study that was approved by the Ethics Committee of the General University Hospital in Prague, Czech Republic, in accordance with the Declaration of Helsinki.

**Table S1** Subject group characteristics for the PD cohort

| Group | PD patients | Healthy controls |
|---|---|---|
| **N** | 30 | 30 |
| **Age (mean±SD [range])** | 64.6±7.7 [46-82] | 63.5±7.9 [46-83] |
| **sex (male/female)** | 13/17 | 15/15 |
| **UPDRS total off (mean±SD [range])** | 45.3±15.0 [11-95] | - |
| **UPDRS total on (mean±SD [range])** | 23.9±10.6 [5-47] | - |

UPDRS – Unified Parkison's Disease Rating Scale, SD – standard deviation

##### *Image pre-processing of the PD dataset*

Pre-processing of all data was conducted in Matlab using SPM12 (Friston et al., 1994). Pre-processing of the rsfMRI data from the PD dataset comprised co-registration of functional and structural images, normalization into Montreal Neurological Institute (MNI) space based on structural information, masking of non-grey matter voxels and smoothing with Gaussian kernel of 6 mm FWHM. Motion (24 parameters based on Friston-24, Friston et al., 1996) was regressed out aside with mean white matter and cerebrospinal fluid signals. Fractional Amplitude of Low Frequency Fluctuations (fALFF) was computed as a measurement of local activity using the REST toolbox with default parameters (linear detrending and 0.01-0.08 Hz band-pass filtering).

**Risperidone cohort**

The risperidone study was conducted using a double-blind, randomized, three-period (each one week apart) cross-over design. Twenty-one healthy volunteers underwent three imaging sessions following single dose administration of either low or high dose of risperidone or placebo (Table S2).

**Table S2** Subject group characteristics for the risperidone cohort

| | **Risperidone cohort** |
|---|---|
| **N** | 21 |
| **Demographics (n male, age±SD)** | 21, 28±7 |

SD – standard deviation

**MRI data and image processing for the PD cohort**

Imaging was performed using a 3T MR scanner (Siemens Skyra) with a 32-channel head coil. During rsMRI scanning, 304 whole brain image volumes consisting of 30 slices were continuously acquired using a standard T2*-weighted echo-planar imaging sequence (TR=2s, TE=30ms, voxel size = 3*3*3 $mm^3$). Additionally, for all subjects a standard T1-weighted magnetisation-prepared rapid gradient echo (MPRAGE) image scan with 1 mm isotropic resolution was acquired for structural co-registration purposes.

All image processing was performed using Statistic Parametric Mapping (SPM12) software and the REST toolbox [Friston et al., 1994; Song et al., 2011]. Pre-processing comprised removal of the first 4 dummy scans, anatomical motion and distortion correction, co-registration to the structural MRI, normalization to the Montreal Neurological Institute (MNI) space using information derived from the structural scan, masking of non-grey matter voxels and smoothing with a Gaussian kernel of 6 mm full width at half maximum (FWHM). Further pre-processing performed in the REST toolbox prior to fALFF computation comprised removal of a linear trend and bandpass filtering using default parameters (0.01-0.08 Hz).

##### MRI data and image processing for the risperidone cohort

All scans were conducted on a GE MR750 3Tesla scanner using a 12-channel head coil. ASL image data were acquired using a pseudo-continuous Arterial Spin Labelling sequence (PCASL) with a multi-shot, segmented 3D stack of axial spirals (8-arms) readout with a resultant spatial resolution of 2x2x3mm. Three control-label pairs were used to derive a perfusion weighted difference image [Dai et al., 2008]. The labelling RF pulse had a duration of 1.5s and a post-labelling delay of 1.5s. The sequence included background suppression for optimum reduction of the static tissue signal. A proton density image was acquired in 48sec using the same acquisition parameters in order to compute the CBF map in standard physiological units (ml blood/100gm tissue/min).

Pre-processing of CBF data was performed using the Statistical Parametric Mapping (SPM12) software package [Friston et al., 1994]. CBF data were first co-registered to individual structural T1 images. Structural scans were segmented and normalized into the Montreal Neurological Institute (MNI) space using the SPM Segment function. Deformation parameters derived from this normalization were then applied to the co-registered CBF data to bring them into the MNI space.
